# Neural signatures of visual awareness independent of post-perceptual processing

**DOI:** 10.1101/2023.06.09.543951

**Authors:** Michael A. Cohen, Cole Dembski, Kevin Ortego, Clay Steinhibler, Michael Pitts

## Abstract

What are the neural processes associated with perceptual awareness that are distinct from pre-conscious sensory encoding and post-perceptual processes such as reporting an experience? Here, we used electroencephalography (EEG) and a no-report visual masking paradigm with parametric manipulations of stimulus visibility to search for neural signatures associated with perceptual awareness independent from both early sensory processing and subsequent reporting. Specifically, we manipulated the time between stimuli and subsequent masks in a linear manner (i.e., 17ms, 33ms, 50ms, 67ms, and 83ms) such that observers’ awareness of the stimuli increased in a *non-linear* fashion (i.e., stimuli were never seen at the two shortest time intervals, always seen at the two longest intervals, and seen approximately 50% of the time at the intermediate interval). Moreover, we manipulated the task across blocks to create separate report and no-report conditions. Overall, we found one neural signal that was closely associated with perceptual awareness, independent from the task: a fronto-central event-related potential (ERP), from ∼250-300ms, that we refer to as the N2. In contrast, earlier ERP signals were linked with the linear manipulation of stimulus strength regardless of visibility, while later candidate signatures, such as P3b and temporal generalization of decoding, were present in the report condition but absent in the no-report condition suggesting a closer association with task performance than perceptual awareness. Taken together, these findings inform current debates regarding theories of consciousness and offer new avenues for exploring the neural mechanisms supporting conscious processing.

**Significance statement:** What differentiates conscious and unconscious processing in the brain? Here, we identify an electrophysiological signature of perceptual awareness using a combination of visual psychophysics and electroencephalography (EEG). In addition, we used a newly developed “no-report” paradigm, in which observers did not report anything about their perceptual experience to separate neural signals associated with consciousness from those associated with the act of reporting (i.e., memory, motor planning, etc.). Using this no-report paradigm was critical because several other candidate signatures of conscious processing were present when observers reported their experiences but completely disappeared when observers did not report their experiences. These findings open the door to future research interested in the neural mechanisms associated of conscious processing.

## Introduction

What neural processes differentiate conscious and unconscious processing? To answer this question, researchers often compare neural responses when observers report being aware vs. unaware of a stimulus (Dehaene, 2014). However, results stemming from this approach are largely united by an inconspicuous flaw. Whenever observers are aware of a stimulus, they are asked to report whether they detected it or to report something about what they perceived. However, such judgements cannot be made when a stimulus goes unnoticed. Therefore, in report-based paradigms, certain post-perceptual processing steps occur only when observers perceive a stimulus, leading to an overestimation of the neural mechanisms supporting conscious processing (Tsuchiya et al., 2015).

This insight led researchers to develop “no-report” paradigms in which observers do not provide reports about their perceptual experience (Pitts et al., 2012; Frässle et al., 2014; Kapoor et al., 2022). The goal of no-report paradigms is to minimize the effect of post-perceptual processing on the neural activity associated with perceptual awareness. Indeed, numerous studies have shown that candidate signatures of conscious processing, such as the P3b or fronto-parietal activation, disappear in no-report paradigms (Cohen et al., 2020; Hatamimajoumerd et al., 2022).

If the P3b and fronto-parietal activation are not signatures of conscious processing, what are? To answer this question, we combined the no-report methodology with a parametric manipulation of stimulus strength. Specifically, we manipulated stimulus visibility by linearly increasing the stimulus onset asynchrony (SOA) between face stimuli and subsequent masks (i.e., evenly-spaced SOAs of 17ms, 33ms, 50ms, 67ms, and 83ms). Critically, when the time between the faces and mask increases linearly (i.e., a stimulus-based difference), the rate at which stimuli are consciously perceived (i.e., a perception-based difference) increases in a non-linear, bifurcated fashion (Del Cul et al., 2007). Any neural signal that displays a non-linear, bifurcated pattern across SOAs is a potential correlate of conscious processing, while signals showing non-bifurcated (e.g., linear) patterns across SOAs can be ruled-out as candidates. Furthermore, by performing this experiment under conditions of no-report, we can also rule-out any bifurcated signals linked with post-perceptual processing, leaving only those associated with perceptual awareness. ^1^ Although the no-report methodology and linear/non-linear manipulations have been employed in previous work (Del Cul et al., 2007; Kouider et al., 2013; Sergent et al., 2021) this study is the first instance of these two approaches being brought together in the domain of visual awareness. Overall, the main result that would link a neural signal to perceptual awareness would be finding that in the no-report condition, a neural signal mirrored the bifurcated pattern of behavioral responses from the report condition.

To preview the results, we identified a neural signal that matched the behavioral results in this critical way: an ERP from ∼250-300ms measured over a set of fronto-central electrodes, which we refer to as the N2. In the no-report condition, the N2 increased in a non-linear, bifurcated manner that mirrored the psychometric function of behavioral responses from the report condition. This N2 stands in contrast to the P1, which increased in a non-bifurcated (roughly linear) fashion across SOAs. Meanwhile, the P3b (Dehaene et al., 2014) and late, sustained temporal generalization of decoding (King et al., 2014) disappeared under conditions of no-report. Finally, an earlier signal (N170/VAN) failed to show bifurcation dynamics. These findings suggest that the neural processes associated with perceptual awareness independent of post-perceptual processing arise earlier than predicted by cognitive theories of consciousness (Odegaard et al., 2017; Dehaene et al., 2017), but later than predicted by sensory theories of consciousness (Koch et al., 2016; Lamme, 2018).

## Methods

### Pre-registration and data sharing

The methods and planned analyses were pre-registered on the Open Science Framework (OSF): https://osf.io/6d5av/. Time windows and electrodes for the ERP analyses of the P1, N2, and P3b were established based on data from an initial set of 15 participants and included in the pre-registration before data acquisition began (see ERP Analysis, below). None of these 15 participants were included in any of the final ERP analyses reported below. Raw and processed EEG data, behavioral data, stimuli, and experiment codes are also available on OSF.

### Participants

56 participants between the ages of 18-29 participated in the experiment. Data from 15 participants were used to define the time-windows and electrodes for ERP analyses. 8 participants were unusable due to technical issues during EEG recording, and 8 participants were excluded for failing to meet the established inclusion criteria (see Exclusion Criteria, below), resulting in a total of 25 participants in the final sample. All participants had normal or corrected-to-normal vision, no known neurological conditions, and no concussions in the preceding year. The experimental procedure was approved by the Institutional Review Board at Reed College and informed consent was obtained from each participant before the experiment, who were compensated for their time.

### Stimuli

Stimuli consisted of 40 grayscale face images (120 x 140 px; 2.7° x 3.15°) and 40 masks embedded in a black background adapted from Kouider et al. (2013). Masks were composed of scrambled, inverted faces and oval objects, and were similar to the face stimuli in size and shape. The contrast level of the face stimuli was adjusted to match each participant’s perceptual threshold as determined by a staircase procedure completed prior to beginning the main experiment (see below).

The green rings (RGB 0, 255, 0) that served as targets in the no-report condition were 15 px wide with a horizontal diameter of 120 px and a vertical diameter of 140 px, measured from outer edge to outer edge. These dimensions ensured each face was fully encircled and minimized the negative space between the stimuli and ring without occluding facial features.

Stimuli were presented centrally against a black background at 120 Hz on a 1920 x 1080 px BenQ monitor, positioned at eye level approximately 70 cm in front of the seated participant. A small red fixation dot (RGB 255, 0, 0; 5 x 5 px; 0.23°) was centered on the screen throughout trial presentation. Neurobehavioral Systems’ Presentation 19.0 was used to present the stimuli in the main experiment, and stimuli were controlled using Psychtoolbox 3.0.14 (Kleiner et al., 2007) for MATLAB (v. 2017a) during the thresholding procedure.

### Experimental design

Broadly speaking, our masking paradigm was modeled after the experiments by Del Cul et al. (2007) and Kouider et al. (2013). Each participant first completed a staircase procedure to determine their perceptual threshold (see below), followed by a brief demonstration of the stimuli, and then the main experiment. The primary experimental procedure consisted of 32 blocks of 60 trials, divided into two 16-block sections with one section being the report condition and another being the no-report condition. All participants completed both the report and the no-report conditions and the order of the conditions was counterbalanced across participants. Participants were given written instructions on the screen at the start of each condition.

Each block began with the appearance of a central red fixation dot that remained on-screen throughout the block. Participants were instructed to keep their eyes on the dot and avoid eye movements and excessive blinking. On trials in which a face was present, a face stimulus was presented in the middle of the screen for 8.33ms, then followed by a blank screen for a variable delay, then followed by two consecutive masks shown for 100ms, and then finally a randomly jittered inter-trial interval of 1050-1450ms, which served as a response window in the report condition (Figure 1A).

**Figure 1.**
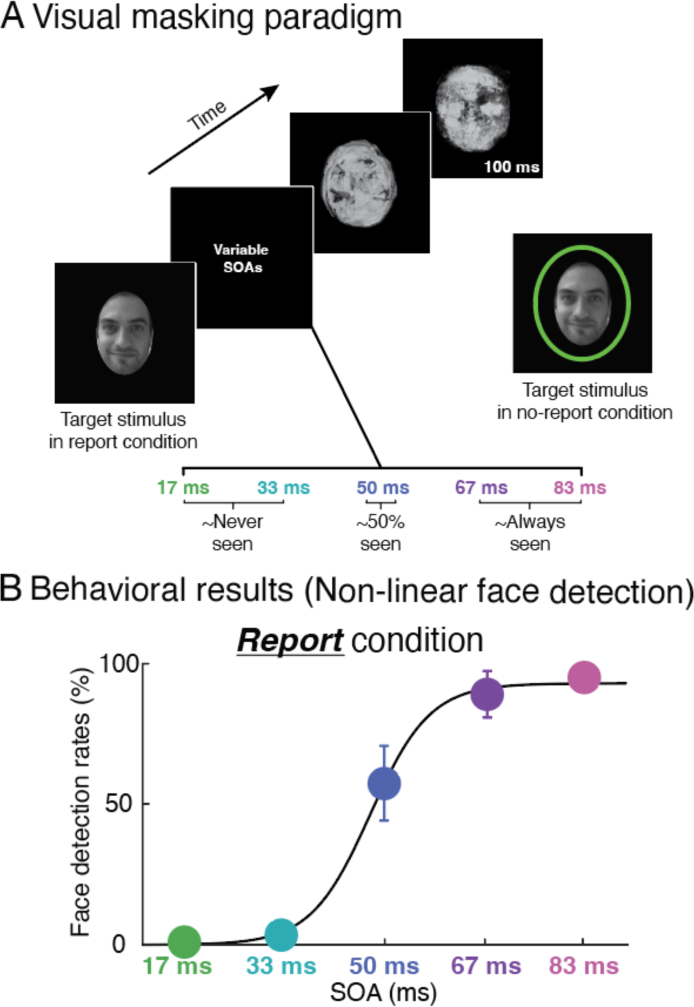
Behavioral paradigm and results. A) Design of visual masking paradigm. A face (or blank) was presented for 8ms, followed by a variable SOA, and then two successive masks, each presented for 100ms. In the report condition, subjects were tasked with detecting faces on each trial (Y/N). In the no-report condition, the subjects’ task was to detect occasional green ring targets randomly presented throughout the stream (i.e., sometimes superimposed on a face and sometimes shown when no face is present). B) Report condition behavioral results. Face detection rates are presented on the y-axis and the different SOAs are presented on the x-axis. Error bars represent the standard error of the mean and the five data points are fit with a non-linear regression.

On each trial, the stimulus-onset asynchrony (SOA) between the face stimulus and the first mask could be 16.67ms, 33.33ms, 50ms, 66.67ms, or 83.33ms (SOAs ‘17,’ ‘33,’ ‘50,’ ‘67,’ and ‘83,’ respectively). The SOA on any given trial was selected randomly, but all SOAs appeared the same number of times in a block. These SOA durations were chosen after pilot testing demonstrated that, when stimulus contrast was adjusted according to each participants’ perceptual threshold so that they could detect the stimulus half of the time on SOA 50 trials, participants were almost never able to see the stimuli on SOA 17 and 33 trials, and almost always saw them on SOA 67 and 83 trials. Linearly manipulating the SOA duration across these five intervals led to a clear sigmoidal psychophysical function, with minimal differences in visibility between the two shortest SOAs (17 and 33ms) or the two longest (67 and 83ms), and maximal differences around the threshold SOA (33 vs. 50ms and 50 vs. 67ms).

On 17% of trials, no face stimulus was presented at all. This provided a set of ‘mask-only’ trials that were used in analysis to isolate stimulus-related from mask-related brain activity (see ERP Analysis, below). Independently, on 15% of trials a green elliptical ring appeared with the same timing as the face stimulus, centrally located and sized so as to encircle the same area where the face stimuli were presented (‘green ring’ trials). These were used as targets in the no-report condition and excluded from all further analyses.

In the report condition, participants indicated via button press after every trial if they had or had not seen a face. These results were used to confirm that the SOA manipulation successfully led to bifurcated perceptual reports, and as a replication of Del Cul et al. (2007).

For the no-report condition, participants were instructed to press a button whenever they saw a green ring that randomly appeared in roughly the same location as where the face stimuli would be shown. Importantly, the green ring did not only appear superimposed around a face; rather, it was presented at the same rate (15%) on trials when no face was shown. As a result, the face stimuli were spatially and temporally attended but were task-irrelevant and were not followed by a response. The task was designed to be challenging enough to keep the participant engaged but not so demanding that they might become intentionally blind to the task-irrelevant face stimuli.

After each 60-trial block, participants received feedback on their task performance. In the report condition, they were given the percentage of all faces they had detected and their false alarm rate (i.e., if they responded “seen” to any mask-only trials). Since participants were not supposed to see any of the faces at the shortest SOAs (and only half the trials at SOA 50) but those trials still counted as “misses” for the purposes of scoring, participants were informed that the expected detection rate was around 50%. This was so participants did not become frustrated with the task due to perceived low-performance. In other words, if participants believed they were expected to see 100% of the faces but were unable to score above 50%, they would be more likely to feel discouraged and lose engagement with the task or alternatively adjust their response criterion which would increase the rate of false alarms.

Feedback in the no-report condition consisted of a count of the number of rings they had missed and it was emphasized to the participants that they should not be missing any of the rings. However, in pilot testing, we found that some participants were unable to see the green rings at all on SOA 17 trials, thus these most difficult trials were not counted as misses. Participants were informed of this at the start.

The experiment began with a staircase procedure manipulating the contrast of the face stimuli to establish each participant’s perceptual threshold (QUEST: Watson et al., 1983). The trial design and task were almost identical to the report condition of the main experiment except all trials used a stimulus-mask SOA of 50ms and no green rings were presented. Additionally, the program paused after each trial to wait for the participant’s response. The results of this procedure were used to set the contrast of the stimuli for the rest of the experiment for each individual participant.

This was followed by a short ‘training’ session to further familiarize participants with the stimuli and show them the green rings. Participants were shown 25 trials at each SOA, in order from longest to shortest. The last five trials in the sets were always green ring trials, and participants were given the option of re-watching each set before going on to the next.

### EEG acquisition

Scalp EEG was recorded using custom 64-channel electrode caps with an equidistant M10 layout (EasyCap, Herrsching, Germany). Signals were amplified via two 32-channel Brain Amp Standard amplifiers (Brain Products, Gilching, Germany), online filtered from 0.1 to 150 Hz, and digitized at 500 Hz. All channels were referenced to an electrode on the right mastoid during recording and electrode impedance was kept under 10 kΩ. Two horizontal EOG channels positioned laterally to each eye and one vertical EOG channel under the left eye were used to monitor blinks and eye movements.

### Exclusion

Participants were excluded from analysis if over 50% of trials were rejected due to EEG artifacts or if their false alarm rate was higher than 20%. Six participants were excluded due to excessive artifacts and two were excluded for false alarms.

### Data preprocessing

EEG preprocessing was performed in BrainVision Analyzer 2.2 following the steps below:

1. Visual identification of any noisy electrodes and interpolation of surrounding electrodes to reconstruct the noisy channel, if necessary.
2. Re-referencing to the average of the left and right mastoid channels.
3. Bipolar HEOG and VEOG channels constructed by re-referencing the left and right horizontal EOG channels and the vertical EOG channel and electrode 50 (FP1) as bipolar pairs.
4. Segmentation of trials around the stimulus (or the first mask in mask-only trials) and removal of all green-ring trials.
5. Baseline correction using the time window from −200 to 0ms pre-stimulus.
6. Semi-automatic rejection of trials with artifacts using peak-to-peak amplitude thresholds (minimum to maximum) of 50 µV for eye movements and 150 µV for blinks and other artifacts (e.g., electrode drift, muscle noise), with adjustments to these thresholds made as-needed at the participant level.

### ERPs and mask subtraction

Trials were sorted by condition and trial type (SOA or mask only). Each group of trials was averaged at the participant-level and low-pass filtered at 25 Hz. To isolate stimulus-related neural activity from mask-evoked activity, the mask-only ERPs were time-shifted so that the timing of the masks were aligned with mask onset in the stimulus-mask trials (following the same procedure as in Del Cul et al., 2007). This was done for each SOA and these time-shifted mask ERPs were subtracted from the corresponding stimulus-mask ERPs, in the following procedure:

1. The mask-only ERP is initially time-locked to mask onset (i.e. time 0 is when the first mask was presented).
2. Time 0 of the mask-only ERP is shifted by −17, −33, −50, −67, or −83ms to match stimulus onset in each SOA. In other words, the mask-only ERP is shifted forwards in time relative to time 0 so that mask onset is aligned with mask onset in each of the stimulus-mask ERPs.
3. The time-shifted mask ERPs are subtracted from the corresponding stimulus-mask ERPs.

### ERP analysis

#### Electrode and time window selection

Grand-average data from an initial group of 15 participants was used to determine electrodes and time windows for ERP components. Three ERPs of interest were identified based on data from this initial group: the P1, displaying a linear increase in amplitude with SOA in both the report and no-report conditions; the N2, displaying bifurcation dynamics in the no-report condition; and the P3b, displaying bifurcation dynamics in the report condition. Based on the data from these 15 participants, the following time windows and electrodes were selected for each component and pre-registered on OSF (nearest channels of the international 10-20 system reported in parentheses):

P1: 100-140ms, electrodes 41-45 and 53-57 (Iz, Oz, O1, O2, O9, O10, PO7, PO8, PO9, PO10).

N2: 250-290ms, electrodes 1-11 and 17-19 (CPz, CP1, CP2, Cz, C1, C2, C3, C4, FCz, FC3, FC4, Fz, F1, F2).

P3b: 300-500ms, electrodes 1, 4-6, and 13-15 (Pz, P1, P2, CPz, CP1, CP2, Cz).

These time windows and electrodes were used for all analyses of these three ERPs, and tested on an independent group of 25 additional participants to confirm replicability.

The N170/VAN analysis was added post-hoc and measured from 140-200ms at electrodes 53 and 57 (PO9,PO10. Once again, these channels were referenced to an average of the left and right mastoids.

#### Assessment of linearity & bifurcation dynamics

Following the methods of Kouider et al. (2013), ERP linearity and bifurcation was assessed by comparing averaged ERP amplitudes across SOAs. An ERP that scales linearly with stimulus strength will exhibit steady and significant increases in amplitude as SOA increases. In contrast, the amplitude of an ERP displaying bifurcation dynamics will not differ significantly between SOAs 17 and 33 or between SOAs 67 and 83, but there will be a significant increase in amplitude from SOAs 33 to 50 and SOAs 50 to 67.

#### Multivariate pattern analysis

All decoding was conducted using the Amsterdam Decoding and Modeling (ADAM) toolbox (Fahrenfort et al., 2018) for MATLAB. Data was exported from BrainVision Analyzer, formatted for use with the ADAM toolbox using EEGLAB (Delorme et al., 2004) and down sampled to 100Hz. For each SOA and task condition, a linear classifier was trained to distinguish between stimulus-present and mask-only trials using a 10-fold cross-validation procedure. At each timepoint, the decoder was first trained on 9/10ths of the trials and then tested on the remaining subset of trials. This was repeated 10 times such that the decoder was both trained and tested on the entire dataset over the course of the procedure, but the classifier was never tested on data used for training when assessing classifier performance. Trial numbers were balanced across classes so that there were equal numbers of stimulus-present and mask-only trials in each fold. Classifier sensitivity, summarized as the area under the receiver operating curve (AUC), was averaged across the folds to calculate overall classifier performance.

#### Temporal generalization

The temporal generalization analysis technique can be used to characterize neural dynamics of perceptual representations by testing if a classifier trained on data from one point in time can successfully discriminate between classes when applied to data from different timepoints. If a classifier trained at time T1 is able to decode stimulus presence at a level significantly above chance when tested at time T2, it suggests that the neural dynamics supporting above-chance decoding at T1 are recurring at T2 (King et al., 2014). Temporal generalization matrices were computed by training classifiers on one timepoint and testing each at every other timepoint. Classification sensitivity was statistically tested against chance and corrected for multiple comparisons using 10,000-permutation cluster-based testing. In this case, we combined the original 15 participants and the subsequent 25 participants (total of 40 participants) as these analyses were exploratory (i.e., not pre-registered) and do not require selections of spatiotemporal regions-of-interest as in the ERP analyses.

#### Variability analysis

Following the methods of Sergent et al. (2021), the inter-trial variability profiles of EEG amplitudes were analyzed at the participant level for three electrode clusters. Trials were sorted by SOA and condition, and mask-only trials were separated according to condition. Separately for each participant, trial-by-trial EEG amplitudes were averaged across each of the clusters of electrodes used in the ERP analyses. Data were then averaged across 50ms time windows from 0 to 700ms post-stimulus, and the standard deviation across trials of this spatially and temporally averaged EEG signal was calculated for each trial group. To account for individual differences in variability and mask-related effects, the standard deviation of mask-only trials was subtracted from the standard deviation of stimulus-present trials in the same condition at the participant level. These adjusted results were averaged across participants. For this exploratory analysis, we also combined the original 15 participants and the subsequent 25 participants for a total of 40.

## Results

### Behavior

The behavioral results from the report condition are plotted in Figure 1B. Overall, we replicated prior findings showing a non-linear, bifurcated pattern of perceptual reports in which faces were rarely detected at the two shortest SOAs (17ms: 1%; 33ms: 4%), were detected roughly half the time at the middle SOA (50ms: 58%), and were frequently detected at the two longest SOAs (67ms: 89%; 83ms: 95%; Del Cul et al., 2007). This is critical because our EEG analyses explicitly seek to differentiate neural responses that scale in a non-bifurcated manner with SOA duration (i.e., 17ms, 33ms, 50ms, 67ms, and 83ms) from those that bifurcate with perceptual awareness (i.e., behavioral responses in Figure 1B).

To identify potential correlates of perceptual consciousness independent of report, we then looked for neural responses in the no-report condition that matched participants’ bifurcated behavioral responses in the report condition. For a neural signal to bifurcate, it must meet four criteria that we pre-registered in advance: 1) little or no difference between the 17ms and 33ms SOAs, 2) a significant difference between the 33ms and 50ms SOAs, 3) a significant difference between the 50ms and 67ms SOAs, and 4) little or no difference between the 67ms and 83ms SOAs. If any of these benchmarks are not met for a given neural signal in the no-report condition, we would argue that signal does not properly mimic the psychometric function from the behavioral results.

### ERP: P1

Which EEG signals increased in a non-bifurcated manner? In this case, we found that the amplitude of the P1 approximated a linear increase in stimulus-mask timing across the five SOAs (i.e., 17ms, 33ms, 50ms, 67ms, and 83ms, Figure 2). In the no-report condition (Figure 2A), there was a significant increase in the size of the P1 for the first two successive SOA pairings (17 vs. 33ms, *t*(24)= 4.85; *P*<0.001; 33 vs. 50ms, *t*(24)=7.24; *P*<0.001) and smaller (non-significant) increases for the last two SOA pairings (50 vs. 67ms, *t*(24)=.96; *P*=.35; 67 vs. 83, *t*(24)=1.53; *P*=.14). A similar pattern was found in the report condition (Figure 2B): P1 amplitude was significantly different for the first two SOA pairings (17 vs. 33ms, *t*(24)=6.65; *P*<0.001; 33 vs. 50ms, *t*(24)=6.15; *P*<0.001), but not the last two SOA pairings (50 vs. 67ms, *t*(24)=1.46; *P*=.16; 67 vs. 83, *t*(24)=1.88; *P*=0.07). This pattern of non-bifurcated (roughly linear) P1 amplitude change across SOA did not change as a function of the act of reporting: compare Figures 2A and B (interaction between no-report and report: *F*(4,120)=0.78, P=0.54; Table 1).

**Figure 2.**
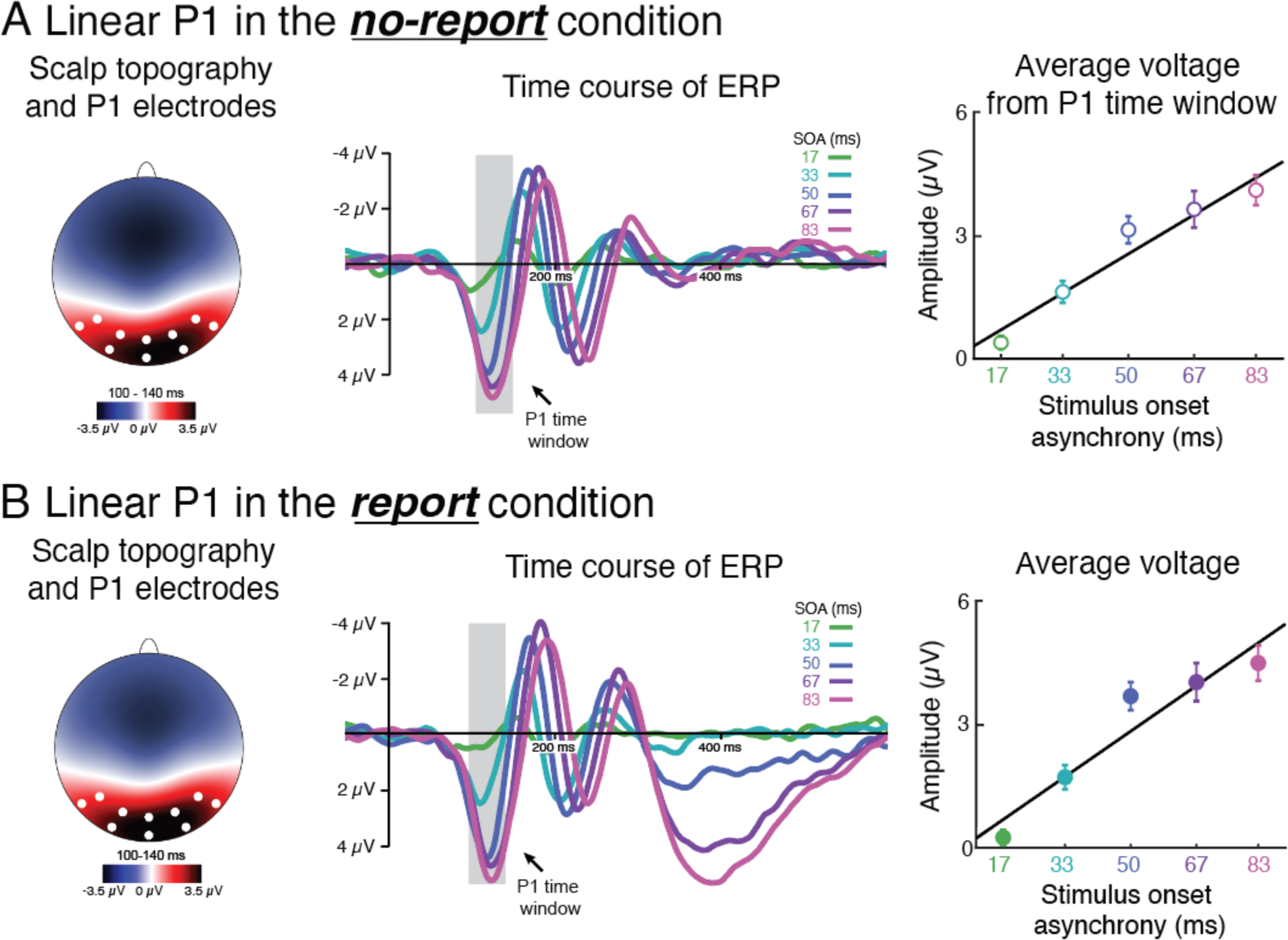
Visualizations of the P1 in the no-report and report conditions. A) No-report and B) report conditions. Left: Topographical voltage distributions between 100-140ms with all electrodes used to analyze the P1 indicated on the scalp map. Center: P1 waveforms for all five SOAs plotted over time. Right: Average amplitudes from electrodes used for the P1 across all five SOAs. Error bars represent the standard error of the mean and the five data points are fit with a linear regression.

### ERP: P3b

The next ERP we examined the P3b. The P3b has been associated with conscious perception across numerous studies (Sergent et al., 2005; Kouider et al., 2013; Dehaene, 2014; Förster et al., 2020; Derda et al., 2019; Ye et al., 2019), and a previous study using the same type of bifurcation logic used here linked perceptual awareness with the P3b under conditions of report (Del Cul et al., 2007). We sought to replicate this result in the report condition and examine whether it would hold in the no-report condition.

The results from the P3b analysis are shown in Figure 3. In the report condition, we observed signs of non-linear bifurcation in the P3b, replicating prior work (Del Cul et al., 2007). There was no difference in P3b amplitude for the two shortest SOAs (17 vs 33ms, *t*(24)=1.76; *P*=.86; Table 1), along with significant increases in amplitude surrounding the middle SOA (33 vs. 50ms and 50 vs 67ms; t(24)>5.77; *P*<0.004), consistent with bifurcation dynamics. However, we also found a significant increase in P3b amplitude between the two longest SOAs (67 vs 83ms; (*t(24)*=7.07; *P*<0.001). Most importantly, we found no evidence of a P3b in the no-report condition, during which mean amplitudes never rose significantly above zero at any SOA (*t(24)*>1.95; *P*>0.06 in all cases; Figure 3A; interaction between no-report and report P3b amplitudes: *F(*4,120)=52.19, *P*<0.001). The complete disappearance of these effects in the no-report condition reinforces findings from prior work showing that the P3b is associated with task performance, such as reporting a perceptual experience, rather than with the experience itself (Pitts et al., 2012; Pitts et al., 2014; Shafto and Pitts, 2015; Cohen et al., 2020; Chen et al., 2022).

**Figure 3.**
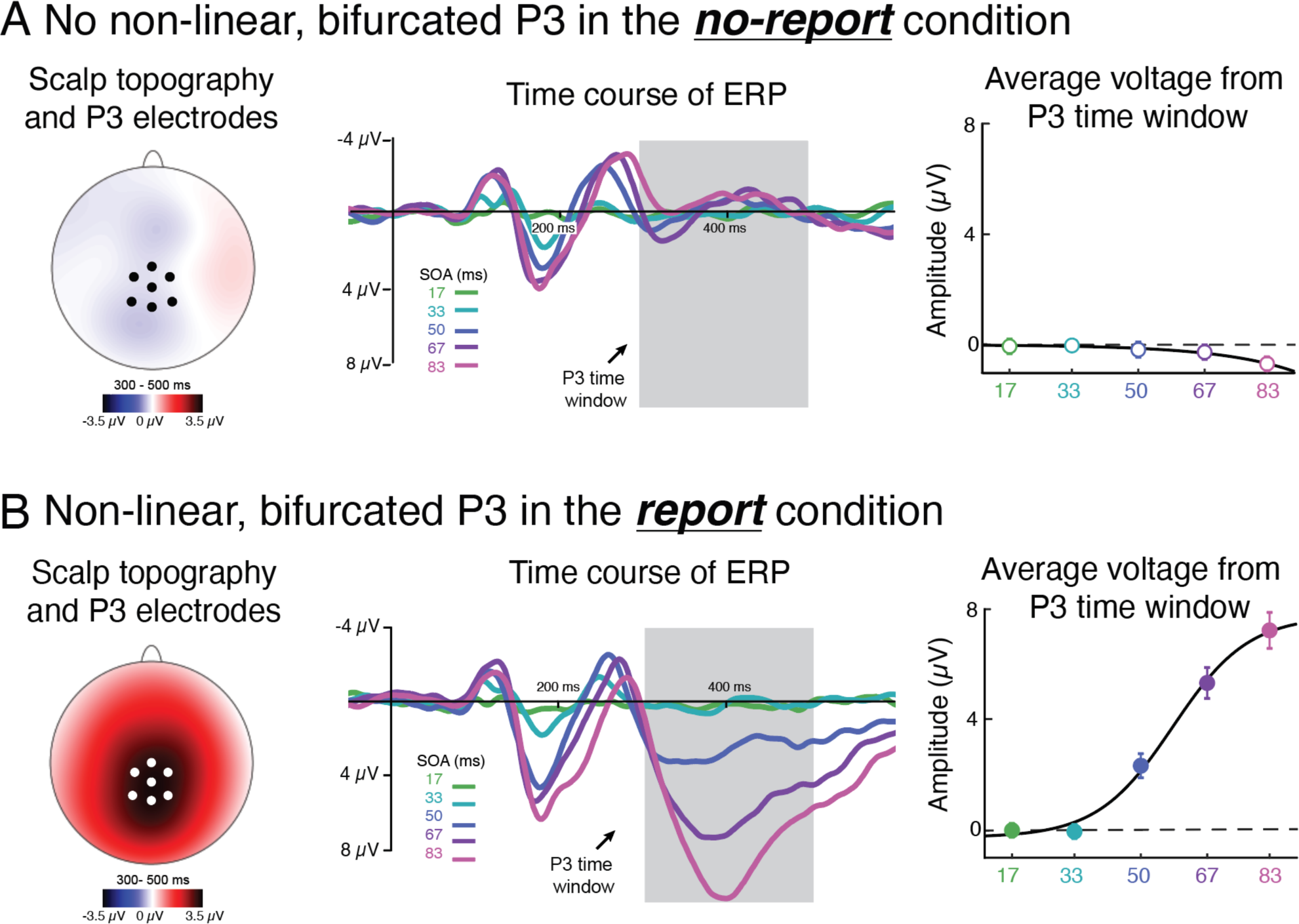
Visualizations of the P3b in the no-report and report conditions. A) No-report and B) report conditions. Left: Topographical voltage distributions between 300-500ms with all electrodes used to analyze the P3b indicated on the scalp map. Center: P3b waveforms for all five SOAs plotted over time. Right are the average amplitudes from all electrodes used for the P3b across all five SOAs. Error bars represent the standard error of the mean, and the five data points are fit with a non-linear regression.

### ERP: N170/VAN

After ruling out the P1 and P3b, we looked next at negative-going waveforms between the P1 and P3b time-windows, which have been previously proposed as potential neural correlates of perceptual awareness (Dembski et al., 2021). For example, the “visual awareness negativity” (VAN) has been defined as a relatively more negative ERP signal for seen compared to unseen stimuli, with a bilateral posterior distribution, roughly in the time window between 150-300ms (Förster et al., 2020; Dembski et al., 2021). When faces are used as stimuli, it is difficult to distinguish the VAN from the N170, and several studies have proposed the latter as a content-specific correlate of conscious face perception (Reiss et al., 2007; Fisch et al., 2009; Harris et al., 2011; Rodriguez et al., 2012; Navajas et al., 2013; Sandberg et al., 2013; Shafto and Pitts, 2015). In the current analysis, we refer to this effect non-committally as an N170/VAN, and later discuss how future studies might be designed to better distinguish between these highly similar neural signals.

Results from the N170/VAN analysis are shown in Figure 4. In both conditions, the N170/VAN followed a pattern of amplitude increase that was neither linear nor bifurcated. In the report condition (Figure 4B), N170/VAN amplitudes increased significantly across the first three SOA pairings (17 vs. 33ms, *t(*24)=4.48; *P<*0.001; 33 vs. 50ms, *t(24)*=8.68; *P<*0.001; 50 vs. 67ms, *t(24)*=2.38; *P<*0.02; Table 1) and then actually decreased across the two longest SOAs (67 vs. 83ms; t(24)=2.18; P<0.04). Meanwhile, in the no-report condition (Figure 4A), the N170/VAN grew significantly across SOAs until SOA 50 (17 vs. 33ms, *t(*24)=5.17; *P<*0.001; 33 vs. 50ms, *t(*24)=6.80; *P<*0.001), with no further increase in amplitude from SOA 50 to 83 (50 vs. 67ms, *t(24)*=0.05; *P=*0.96); 67 vs. 83ms; t(24)=1.14; P=0.26). There was a significant interaction between the report and no-report conditions due to the amplitude of the N170/VAN increasing for the 50-83ms SOAs in the report condition relative to the no-report condition: *F(*4,120)=5.35, *P*<0.001; Table 1).

**Figure 4.**
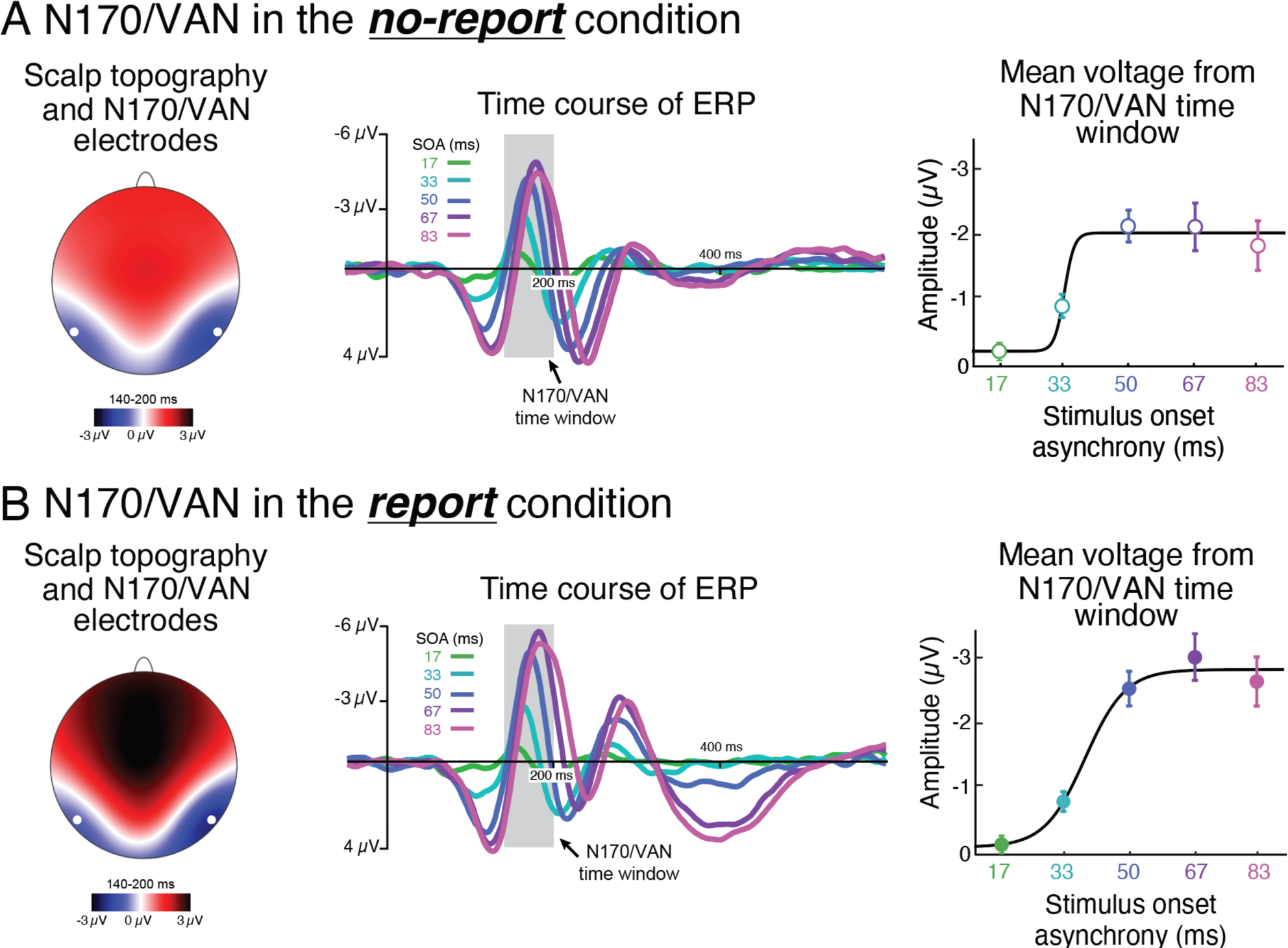
Visualizations of the N170/VAN in the no-report and report conditions. A) No-report and B) report conditions. Left: Topographical voltage distributions between 140-200ms with the electrodes used to analyze the N170/VAN indicated on the scalp map. Center: N170/VAN waveforms for all five SOAs plotted over time. Right: average amplitudes from electrodes used for the N170/VAN analysis across all five SOAs. Error bars represent the standard error of the mean and the five data points are fit with a non-linear regression.

Our findings with the N170/VAN fail to meet our criterion for matching the sigmoid-shaped psychometric function from the behavioral data in a few key ways. First, the amplitude of the signal significantly increased between the 17 and 33ms SOAs. This is inconsistent with the behavioral results since the behavioral difference between these two SOAs was incredibly small (1% vs 3.8%). This finding is in line with previous work reporting an N170 in response to unseen faces (Sterzer et al., 2009; Harris et al., 2013; Suzuki and Noguchi, 2013), and we discuss the implications for the VAN in detail in the discussion below. Second, there is no significant difference between the 50 and 67ms SOAs. This is also inconsistent with the behavioral results since the behavioral difference between these two SOAS was quite large (51.9% vs 86.8%). Taken together, the N170/VAN does not appear to fully track behavioral responses in either condition, suggesting it may not be a candidate signature of perceptual awareness, though more research is called for in the future.

### ERP: N2

In the time-period between the N170/VAN and P3b, one ERP signal closely matched the nonlinear, bifurcated pattern of participants’ perceptual reports: a negativity distributed over fronto-central electrodes which we refer to as the N2 (Figure 5A). Between 250-290ms, this N2 wave in the no-report condition displayed a clear non-linear, bifurcated pattern of amplitude increase that mirrored the psychometric functions in the report condition (compare Figure 1B and Figure 5A right panel). There was no difference in the amplitude of the N2 between the two shortest SOAs: 17ms and 33ms (*t(24)*=0.24; *P*=0.81; Table 1). There were significant differences in N2 amplitudes between 33ms and 50ms (*t(24)*=4.49; *P<*0.001), as well as between 50ms and 67ms (*t(24)*=2.82; *P<*0.01). Finally, N2 amplitudes at the two longest SOAs did not differ significantly: 67ms and 83ms (*t(24)*=0.86; *P*=0.40). This pattern of amplitude changes across SOA met all four of our pre-defined criteria for a bifurcated neural signal.

**Figure 5.**
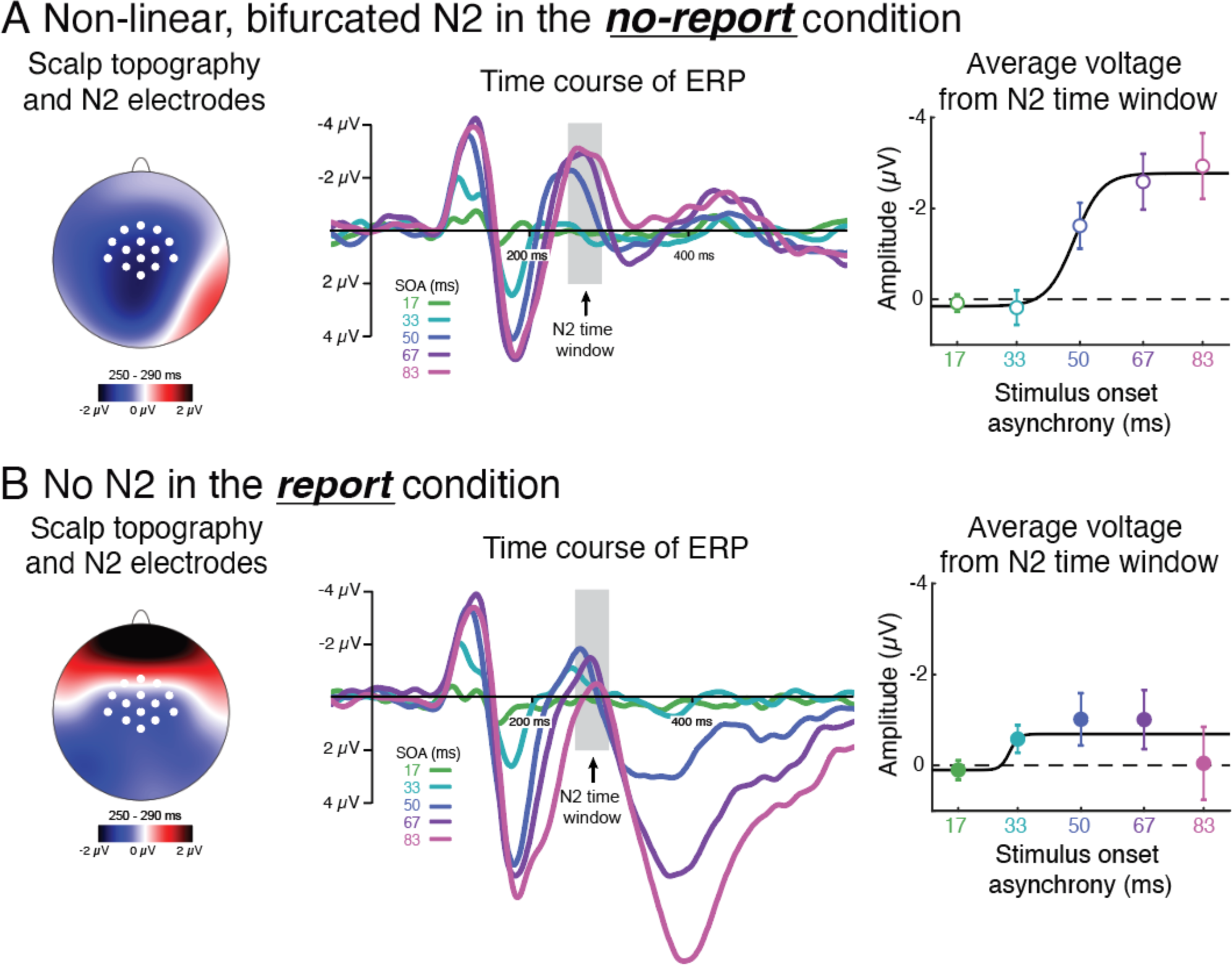
Visualizations of the N2 in the no-report and report conditions. A) No-report and B) report conditions. Left: Topographical voltage distributions between 250-290ms with all electrodes used to analyze the N2 indicated on the scalp map. Center: N2 waveforms for all five SOAs plotted over time. Right: Average amplitudes across all electrodes used for the N2 across all five SOAs. Error bars represent the standard error of the mean and the five data points are fit with a non-linear regression.

It should be emphasized that we measured the N2 in neural responses from the no-report condition, while the perceptual reports it mirrored came from the report condition. As such, the N2 cannot reflect the neural processes underlying the non-linear bifurcation in participants’ perceptual judgments or behavioral responses (i.e., post-perceptual bifurcation dynamics). Instead, the N2 is more likely associated with perceptual awareness per se: in the no-report condition, the stimuli in question were task-irrelevant but otherwise identical to those in the report condition, such that the influence of post-perceptual processing was minimized while the stimuli were presumably perceived at similar rates across SOAs compared to the report condition. Of course, we recognize it cannot be definitively stated that awareness of the face stimuli in the report and no-report conditions was identical. This is an inherent limitation of no-report paradigms that all studies must consider. Here, we specifically designed the no-report task (detection of green rings superimposed on the faces) to minimize differences in attention, effort, etc. across tasks. Nevertheless, we openly acknowledge that we cannot state with full certainty that the rates of face detection were identical in the report and no-report conditions and instead can only make that assumption.

ERP analyses on the same set of electrodes during the report condition revealed a different pattern of results (Figure 5B). The N2 was partially evident, but its pattern of amplitude change across SOAs was clearly affected by the act of reporting the presence of a face stimulus. As a result, the topographical voltage distributions looked quite different between the report and no-report conditions, and none of the bifurcating characteristics of the N2 observed in no-report were noticeably present in the report condition (interaction between no-report and report: *F(*4,120)=11.42, *P<*0.001). There was a significant difference in amplitude between SOAs 17 vs. 33ms (*t(24)*=2.37; *P<*0.05), no difference between SOAs 33 vs 50ms (*t(24)*=0.94; *P*=0.36) and 50 vs. 67ms (*t(24)*=0.02; *P*=0.98), and a significant decrease from SOAs 67 vs. 83 (*t(24)*=2.09; *P*<0.04). None of these patterns are consistent with the trends expected from a non-linear, bifurcated signal. In fact, when comparing the report and no-report conditions directly, there was a significant interaction between the 17 vs. 33ms SOA, the 33 vs. 50ms SOA, and the 50 vs 67ms SOA (*t(24)*>2.19; *P<*0.05 in all cases). This suggests that prior studies may have failed to detect a link between perceptual awareness and the N2 because observers always had to make explicit reports about the contents (or presence) of their perceptual experience.

If the N2 is a true potential signature of perceptual awareness, why was it not found in the report condition? We believe that in the report condition, the spatially and temporally overlapping P3b wave “drowned out” the N2 and prevented it from being detected. To test this possibility, we performed a post-hoc exploratory analysis. In this analysis, we split all 40 participants we ran (i.e., the 15 original and the 25 subsequent) into two groups based on the magnitude of the P3b (i.e., largest P3b group and smallest P3b group; N=20 in each group). We predicted that we would see more “hints” of the N2 amongst the smallest P3b group since the P3b would interfere less with the N2 in that group. Overall, we found no hints of the N2 amongst the largest P3b group since the shape of the N2 function did not match the sigmoid-shaped psychometric function from the behavioral data (Figure 6A). There was a significant difference in amplitude between SOAs 17 vs. 33ms (*t(24)*=2.14; *P<*0.05), and no differences between any of the other SOA pairings (*t(*24)<1.50; *P*>0.15 in all cases). Meanwhile, we did find some hints of the N2 amongst the smallest P3b group since the shape of the N2 function in this case came closer to matching the shape of the sigmoid-shaped psychometric function from the behavioral data (Figure 6B). However, the statistics did not perfectly match the behavioral data since there was a significant difference between SOAs 17 vs. 33 (*t(24)*=2.54; *P*<0.05). Nonetheless, all other statistical tests did match the predictions of a bifurcated signal (SOAs 33 vs. 50: *t(*24)=2.23; *P*<0.04; SOAs 50 vs. 67 *t(*24)=2.07; *P*<0.05; SOAs 67 vs. 83: *t(*24)=0.34; *P*=0.74). While this analysis is by no means definitive, it suggests that interference from the P3b in the report condition may have prevented us from observing an N2 in that condition. Moreover, we believe this finding highlights the importance of no-report paradigms since the neural processes most closely associated with perceptual awareness may simply go unnoticed due to interference from the overlapping neural activity supporting the act of reporting.

**Figure 6.**
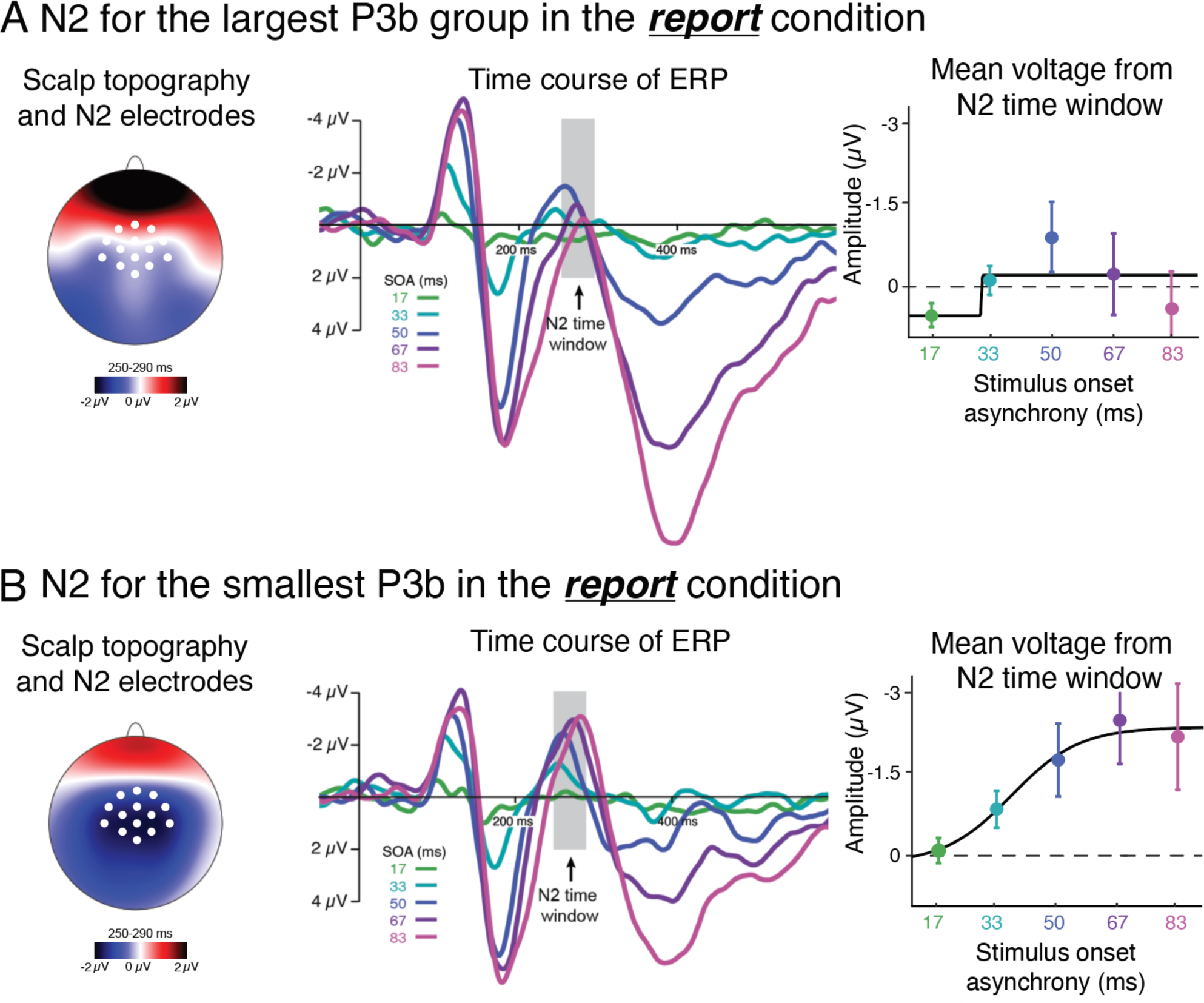
N2 waveforms in the report condition from two groups of participants sorted by P3b peak amplitude (median split, N=20 in each group). A) Waveforms for the largest P3b group and B) waveforms for the smallest P3b group. Left: Topographical voltage distributions between 250-290ms with the electrodes used to analyze the N2 indicated on the scalp map. Center: N2 waveforms for all five SOAs plotted over time. Right: average amplitudes from electrodes used for the N2 analysis across all five SOAs. Error bars represent the standard error of the mean and the five data points are fit with a non-linear regression.

### Temporal generalization of decoding

Having found the N2 to be the only ERP that clearly follows a bifurcated response pattern that reflects perception in the no-report condition, we proceeded to ask if any other EEG signals might display these non-linear dynamics. To examine this possibility, we performed an exploratory analysis on another posited signature of conscious processing, the presence of late metastable activity (Bekinschtein, et al., 2009; King and Dehaene, 2013; Dehaene, 2014; Schurger et al., 2015; Marti and Dehaene, 2017; Derda et al., 2019). Under this proposal, as information processing transitions from unconscious sensory stages to conscious perception, it is stabilized for a few hundred milliseconds so that conscious content may be used to achieve specific goals (i.e., decision-making, verbal responses, motor outputs, etc.). This metastability can be represented in temporal generalization matrices, which visualize how different processing stages and their underlying neural codes unfold over time (King and Dehaene, 2014). Specifically, these matrices are formed by training pattern classifiers to distinguish between two or more stimulus conditions at each time-point and then testing classification performance across all time-points. This process returns a temporal generalization matrix that visualizes the decoding accuracy for every pair of training and testing time points. Several studies have found that perceptual awareness is linked with the transformation of information into a metastable representation (Schurger et al., 2015; Salti et al., 2015). This late metastability is reflected by an off-diagonal square-shaped pattern in the temporal generalization matrix (black square in right panel of Figure 7).

**Figure 7.**
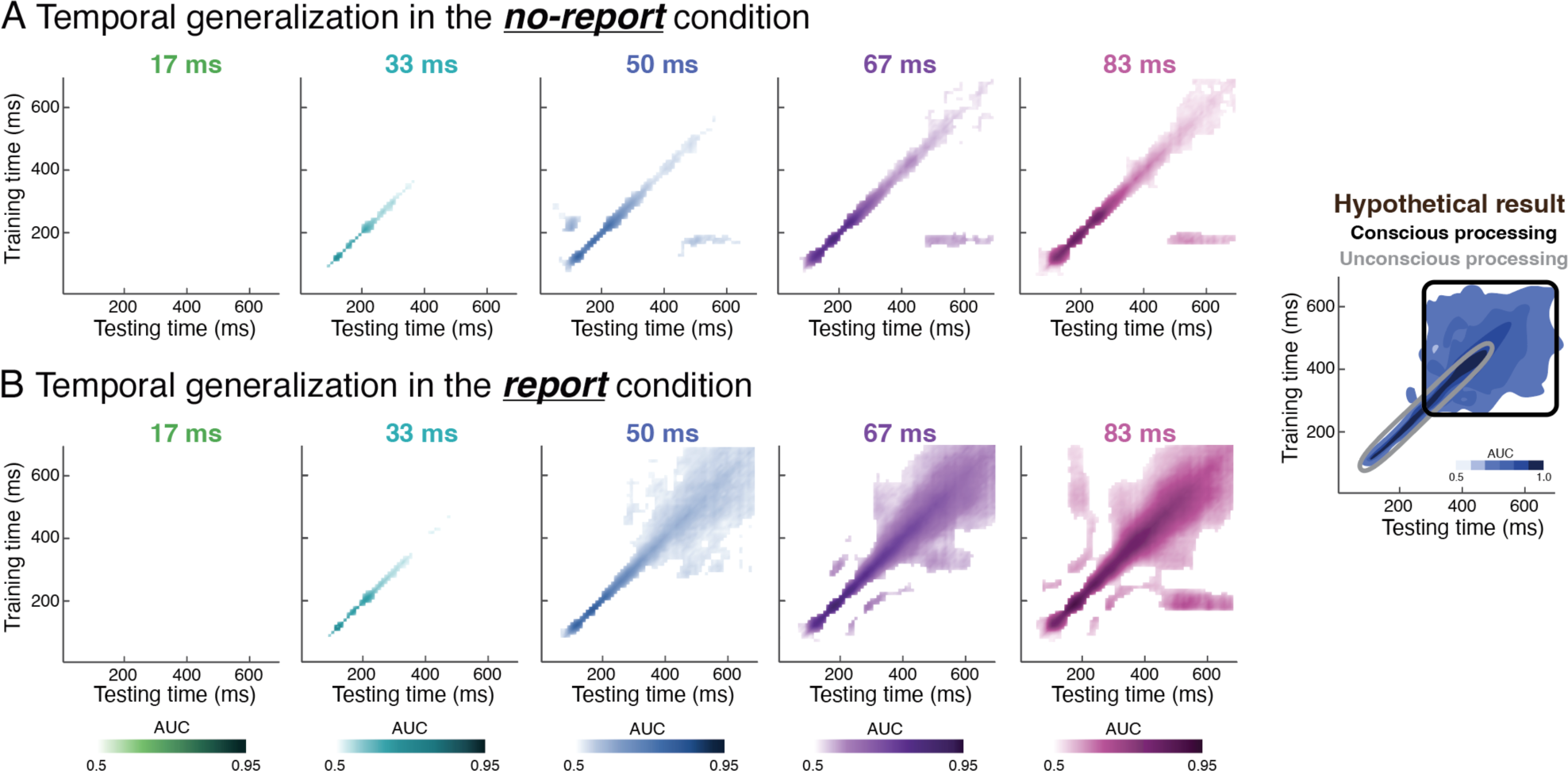
Temporal generalization matrices in no-report and report conditions. A) No-report and B) report conditions. Decoders were trained to discriminate stimulus-present and stimulus-absent trials at each time-point between 0 and 700ms and tested at all time-points. In the no-report condition (A), decoding was successful only along the training-testing diagonal, suggesting a sequential activation of different brain regions. However, in the report condition (B), while the 33ms SOA only allowed for weak decoding along the diagonal, the longer SOAs elicited a combination of diagonal decoding and late sustained generalization (square-shaped patterns) that begins roughly 300ms after stimulus onset. The right panel represents a hypothetical result interpreted under the proposal that late temporal generalization reflects conscious perception, depicting the diagonal associated with unconscious processing and the square-shape associated with conscious processing. The current results challenge this proposal.

To look for evidence of a non-linear bifurcation in this late temporal generalization signal, we trained multivariate decoders to distinguish between trials in which a face was present (stimulus trials) versus absent (mask only trials; see Figure 1) and visualized the results in a standard temporal generalization matrix. In the report condition, we found strong evidence of a metastable representation associated with conscious perception, indicated by late square-shaped patterns in the temporal generalization matrix (Figure 7B). Indeed, at the 50ms SOA, in which observers first started to sometimes notice the face stimuli (see Figure 1B), a late square-shaped pattern emerged that increased in classification accuracy at longer SOAs, when stimuli were more reliably visible. However, in the no-report condition, we found no evidence for a late, sustained, metastable representation: decoding accuracy was only significantly above chance when trained and tested on similar time points (Figure 7A). Together, these results suggest that late, sustained metastability as represented by temporal generalization of decoding is more associated with post-perceptual processing (i.e., reporting tasks) than perceptual awareness per se.

### Inter-trial EEG/ERP variability

Another recently proposed signature of conscious processing independent of report is a spike in inter-trial EEG/ERP variability for stimuli presented at threshold (Sergent et al., 2021). The rationale for this measure revolves around the idea that a threshold stimulus will yield a mixture of high-activity (aware) and low-activity (unaware) trials. This burst in inter-trial variability will not be seen for stimuli that are well above or well below threshold since neural activity should be more homogenous across such trials. Thus, this proposal predicts that we should see an increase in inter-trial variability in the 50ms SOA condition, when the stimulus is at threshold, along with lower variability at the other SOAs which were well below or well above threshold. However, in an exploratory analysis of the current dataset we found minimal evidence of such variability in the no-report condition and no evidence for this variability pattern in the report condition (Figure 8). In the no-report condition, we observed some signs of a peak of inter-trial EEG/ERP variability at the threshold SOA (50ms) during the 250-300ms and 300-350ms time windows, overlapping the time window of the N2 wave (SOAs 33 vs. 50ms: 250-300 ms, *t*(39)=1.76; *P*< 0.05; 300-350ms, *t*(39)=1.68; *P*=0.05), but the difference in variability on SOA 50 vs. 67 trials did not reach significance in either time window (*t*(39)<0.56; *P* >0.32). Inconsistent with previous reports (Sergent et al., 2021), no such pattern was evident during any time window in the report condition.

**Figure 8.**
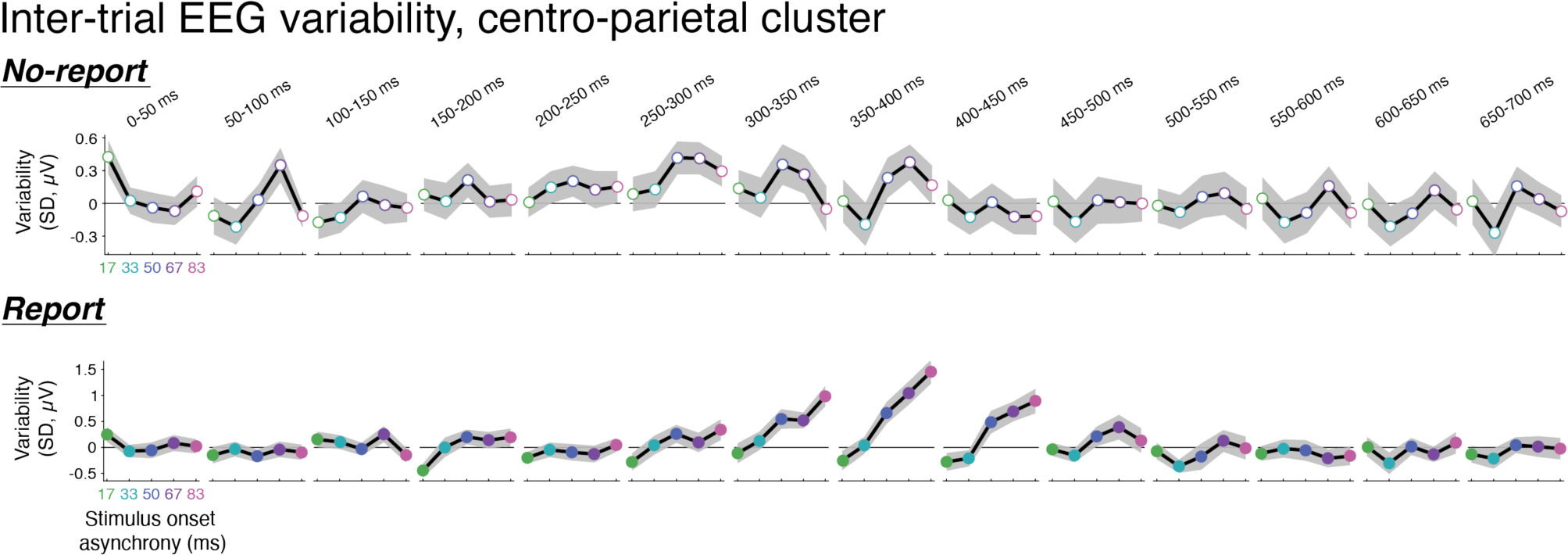
Inter-trial EEG variability in no-report and report conditions at centro-parietal electrodes. Top row corresponds to the no-report condition and the bottom row corresponds to the report condition. On the horizontal axes are the five SOAs (ms) and on the vertical axes is the inter-trial variability of evoked activity as a function of SNR at these different time windows. Note that the electrode cluster used in this analysis is the same one used for the P3b in the present study, which is highly similar to the group of electrodes used by Sergent et al. (35) in their previous variability analysis.

## Discussion

Here, we sought to identify neural signatures associated with perceptual awareness independent of post-perceptual processing. Specifically, we used a novel no-report visual masking paradigm to search for bifurcation dynamics. We found an EEG signal in the no-report condition — the N2 — that displayed bifurcation dynamics closely matching behavioral responses in the report condition. This component differed from the P1 and N170/VAN which showed response patterns inconsistent with bifurcation dynamics, as well as the P3b and late temporal generalization of decoding, which showed bifurcation dynamics in the report condition but disappeared in the no-report condition. Finally, although inter-trial EEG variability analyses showed hints of a peak in variability around the threshold SOA at time-points overlapping the N2 in the no-report condition, this result was inconclusive.

### The N2 and other proposed neural signatures of consciousness

One noteworthy aspect of the N2 is that it ends at ∼300ms, just as the P3b and late temporal generalization begin. Moreover, the initial phase of the P3b appears to overlap the N2 in the report condition, reducing its amplitude and destroying its bifurcation pattern. Thus, an intriguing possibility is that the N2 is associated with perceptual awareness independent of report, and whenever these perceptual experiences are task relevant, this information is then stabilized and fed into other networks associated with maintaining and using those perceptual experiences for behavior. On one interpretation, the N2 might reflect the “global playground” recently proposed by Sergent et al. (2021), while the P3b and late metastability are signatures of the “global workspace” (Mashour et al., 2020). In addition, the timing of the N2 is consistent with a recent study using several neuroimaging techniques along with task irrelevant stimuli to test competing predictions of global workspace and integrated information theories of consciousness (Cogitate et al., 2023).

If the P3b is not related to conscious processing, what is it related to? Across several decades, many potential answers to this question have been proposed (Polich, 2020). Some of the leading hypotheses include: reactivation of stimulus-response links, memory storage, closure of cognitive epochs, and decision processes (Verleger, 2020). Another suggestion is that the P3b is linked with conscious experience, but only when the task requires that experience to be cognitively accessed for report or further processing (Naccache, 2018; Tagliabue et al., 2019; Sergent, 2021). However, in standard “oddball” tasks in which participants consciously perceive “standard” stimuli (e.g., presented on 90% of trials) and “deviant” stimuli (e.g., presented on 10% of trials), the P3b is robust for deviant stimuli but absent for standard stimuli even when both standard and deviant stimuli must be reported (Squires et al., 1975; Duncan-Johnson & Donchin, 1977). While our study used an “exclusionary approach” (Passler, 2023) to rule-out a close relationship between the P3b and visual awareness, it will be important for future work to uncover what precisely the P3b is directly related to.

Recently, Sergent and colleagues (2021) suggested that inter-trial EEG/ERP variability may be a signature of conscious processing independent of report. One of their main findings was a burst of inter-trial variability emerging for stimuli presented around the threshold of consciousness in both report and no-report conditions. This variability showed bifurcation dynamics that matched behavioral responses in the report condition. Strikingly, the time window of this bifurcated variability was almost identical to the time window of the N2 (∼250-300ms). Interestingly, although Sergent and colleagues’ study and the present study examined different sensory modalities, manipulated perceptual awareness in different ways, and used different measurements, the findings from both studies converge on a timeframe of neural dynamics linked with perceptual awareness. Together, these results form a new hypothesis that the neural mechanisms supporting conscious perception begin ∼250-300ms after stimulus onset, during a brief stage of processing in-between low-level sensory encoding and high-level broadcasting across global networks.

Although this study did not entirely replicate Sergent et al.’s (2021) results, we did observe some signs of inter-trial variability between 250-350ms in the no-report condition. While it is unclear why there was no trace of this pattern in the report condition, several viable explanations exist, including the different stimulus modality and methods used to render stimuli invisible. In line with the latter explanation, the current results appeared to show a systematic increase in inter-trial EEG/ERP variability across SOA during most of the P3b time window (300-450ms), which was not observed in Sergent and colleagues’ data. Such an increase in variability could be explained by the additional cognitive demands of perceiving two competing stimuli at longer SOAs (stimulus and mask) compared to perceiving one stimulus at shorter SOAs (the mask). Such differential cognitive demands across SOAs would only be expected to come into play in conditions in which the stimulus is task-relevant.

Another potential neural signature of visual consciousness is the VAN (Koivisto and Revonsuo, 2007; Förster et al., 2020; Dembski et al., 2021). Our data, however, suggest that the N170/VAN may not be directly linked with perceptual awareness because the amplitude across SOAs did not match the behavioral responses in two key ways. First, the N170/VAN was significantly different between 17ms and 33ms while there was very little difference in behavioral responses at these time windows. Second, there was no difference between the N170/VAN at 50ms and 67ms even though there was a significant difference in the behavioral responses at these time windows. This pattern of results differs from the N2, which closely matched the pattern of behavioral responses. It should be noted that some might consider what we refer to as the N2 as the VAN since it is a negative-going waveform occurring within the VAN time window (∼150-300ms). While this is true, the N2 has a spatial distribution that is more anterior than is typically observed for the VAN (Förster et al., 2020). That being said, it remains an open question whether the scalp distribution of the VAN changes for different contents of consciousness within a given sensory modality or for different manipulations of awareness (Dembski et al., 2021).

### The N2 and theories of consciousness

How do these findings relate to the longstanding debate between “cognitive” and “sensory” theories of consciousness? Cognitive theories like the Global Neuronal Workspace or Higher-Order Thought Theory (Dehaene, 2014; Odegaard et al., 2017; Brown et al., 2019; Mashour et al., 2020), are often associated with the P3b and sustained temporal generalization of decoding. Meanwhile, sensory theories like Recurrent Processing and Integrated Information Theory (Koch et al., 2016; Boly et al., 2017; Lamme, 2018), are often associated with earlier electrophysiological signatures like the VAN.

The current results do not fit neatly into either of these frameworks. In terms of both time and space, the N2 appears to index a stage of processing in between early recurrent processing and late global “ignition”. Two hallmarks of the P3b are its sustained response and its broad spatial distribution. These features stand in contrast to the N2, which only lasts 50-100ms, and is spatially circumscribed with bilateral posterior/inferior positive foci (perhaps the other end of its dipole generators) distinct from the P3b. Meanwhile, compared to the VAN, the N2 scalp topography is more anterior, and its timing is rather late to be considered a marker of early recurrent processing. Thus, the N2 appears to be situated in an intermediate stage of processing between the neural markers most often cited as supporting these two classes of theories. Interestingly, a classic study of visual awareness (Sergent et al., 2005) appears to have identified this same signal (see Figure 3 & 7; component labeled N2-N3), but at the time, prior to the advent of no-report paradigms, the authors chose to focus on the larger P3b signal.

Identifying where the N2 originates in the brain will further elucidate how it fits into these frameworks. Unfortunately, we cannot establish its source using scalp EEG alone. However, given its location in space, its timing, and its relation to other established ERPs, the N2 may originate from parietal regions (or interactions between fronto-parietal and occipito-temporal regions). If this proved to be the case, it would not support one of the dominant competing theories over the other. For example, cognitive theorists might see parietal activation as part of a “parietal workspace” in which information reaches consciousness by activating parts of the parietal lobe associated with higher-order functions such as attention, working memory, and decision-making (Emrich et al., 2013; Fedorenko et al., 2013; Woolgar et al., 2018). Conversely, sensory theorists might relate this activity to a posterior cortical “hot zone” that is separate from regions involved in higher-level functions (Koch et al., 2016). Despite the challenges involved in relating such findings to different theories, understanding where the N2 originates will be an important step in the process of triangulating the regions (and interactions between regions) containing the neural networks that support perceptual awareness.

## Acknowledgements

Thanks to Sarah Cormiea, Stanislas Dehaene, Steven Hillyard, Nancy Kanwisher, Ken Nakayama, N. Apurva Ratan Murty, Caroline Robertson, and Claire Sergent for helpful discussions at different stages of this project. This work was supported by National Science Foundation Grant BCS-1829470 to M.A.C. and M.P., Templeton World Charity Foundation Grant TWCF-2022-30267 to M.A.C. and M.P., the Canadian Institute for Advanced Research to M.A.C., the American Psychological Foundation to M.A.C., and the Stillman Drake Fund for Faculty Development to M.P.

## Footnotes

1 While the no-report condition was created to minimize post-perceptual processes, we cannot say with certainty that such post-perceptual processes were entirely eliminated. Indeed, it is possible that with minimal task demands in the no-report condition, participants may nevertheless spontaneously and covertly name, remember, think about, or perform other cognitive tasks on the stimuli. However, this so-called “bored monkey problem” is an inherent limitation of no-report paradigms (Block, 2019) and is not unique to our methodology.

## References

1. Bekinschtein, T.A., Dehaene, S., Rohaut, B., Tadel, F., Cohen, L., & Naccache, L. (2009). Neural signature of the conscious processing of auditory regularities. Proc. Natl. Acad. Sci. USA. 106, 1672–1677.

2. Block, N. (2019). What is wrong with the no-report paradigm and how to fix it. Trends Cogn. Sci. 23, 1003–101.

3. Block, N. (2023) The border between seeing and thinking. Oxford University Press.

4. Bola, M. & Doradzinska, L. (2021) Perceptual awareness negativity—Does it reflect awareness or attention? Front Hum Neurosci. 15, 742513.

5. Boly, M., Massimini, M., Tsuchiya, N., Postle, B.R., Koch, C., & Tononi, G. (2017). Are the neural correlates of consciousness in the front of or in the back of the cerebral cortex? Clinical and neuroimaging evidence. J. Neurosci. 37, 9603–9613.

6. Brown, R., Lau, H., & LeDoux, J.E. (2019) Understanding the higher-order approach to consciousness. Trends Cogn. Sci. 23, 754–768.

7. Chen, Y.K., Cheng, T., & Hsieh, P.J. (2022). P3b does not reflect perceived contrasts. eNeuro. 9.

8. Cohen, M.A., Ortego, K., Kyroudis, A., & Pitts, M.A. (2020). Distinguishing the neural correlates of perceptual awareness and post-perceptual processing. J. Neurosci. 40, 4925–4935.

9. Cogitate Consortium, et al. (2023) An adversarial collaboration to critically evaluate theories of consciousness. bioRxiv. doi: 10.21203/rs.3.rs-310836/v1.

10. Dehaene, S., Naccache, L., Cohen, L., Le Bihan, D., Mangin, J-F., Poline, J-B., & Riviere, D. (2001). Cerebral mechanisms of word masking and unconscious repetition priming. Nat. Neurosci. 4, 752–758.

11. Dehaene, S., Lau, H., & Kouider, S. (2017) What is consciousness, and could machines have it? Science. 358, 486–492.

12. Del Cul, A., Baillet, S., & Dehaene, S. (2007). Brain dynamics underlying the nonlinear threshold for access to consciousness. PLoS Biol. 5, 2408–2423.

13. Delorme, A., & Makeig, S. (2004). EEGLAB: an open source toolbox for analysis of single-trial EEG dynamics including independent component analysis. J. Neurosci. Methods. 134, 9–21.

14. Dembski, C., Koch, C., & Pitts, M.A. (2021). Perceptual awareness negativity: a physiological correlate of sensory consciousness. Trends Cogn. Sci. 25, 660–670.

15. Derda, M., Koculak, M., Windey, B., Gociewicz, K., Wierzchón, M., Cleeremans, A., & Binder, M. (2019). The role of levels of processing in disentangling the ERP signatures of conscious visual processing. Conscious. Cogn. 73, 102767.

16. Duncan-Johnson, C.C. & Donchin, E. (1977) On quantifying surprise: The variation of event-related potentials with subjective probability. Psychophysiol. 14, 456–467.

17. Eklund, R., & Wiens, S. (2019). Auditory awareness negativity is an electrophysiological correlate of awareness in an auditory threshold task. Conscious. Cogn. 71, 70–78.

18. Emrich, S.M., Riggall, A.C., LaRocque, J.J., & Postle, B.R. (2013). Distributed patterns of activity in sensory cortex reflect the precision of multiple items maintained in visual short-term memory. J. Neurosci. 33, 6516–6523.

19. Fahrenfort, J.J., van Driel, J., van Gaal, S. & Olivers, C.N.L. (2018). From ERPs to MVPA using the Amsterdam Decoding and Modeling Toolbox (ADAM). Front. Neurosci. 12, 368.

20. Fedorenko, E., Duncan, J., & Kanwisher, N. (2013). Broad domain generality in focal regions of frontal and parietal cortex. Proc. Natl. Acad. Sci. USA. 110, 16616–16621.

21. Fisch, L., Privman, E., Ramot, M., Harel, M., Nir, Y., Kipervasser, S., Andelman, F., Neufeld, M. Y., Kramer, U., Fried, I., & Malach, R. (2009). Neural “ignition”: enhanced activation linked to perceptual awareness in human ventral stream visual cortex. Neuron. 64, 562–574.

22. Folstein, J.R. & Petten, C.V. (2008). Influence of cognitive control and mismatch on the N2 component of the ERP: A review. Psychophysicology. 45, 152–170

23. Förster, J., Koivisto, M., & Revonsuo, A. (2020). ERP and MEG correlates of visual consciousness: the second decade. Conscious. Cogn. 80, 102917.

24. Frässle, S., Sommer, J., Jansen, A., Naber, M., & Einhäuser, W. (2014). Binocular rivalry: frontal activity relates to introspection and action but not to perception. J. Neurosci. 34, 1738–1747.

25. Hanslmayr, S., Aslan, A., Staudigl, T., Klimesch, W., Herrmann, C.S., & Bäuml, K.H. (2007). Prestimulus oscillations predict visual perception performance between and within subjects. Neuroimage. 37, 1465–1473.

26. Harris, J.A., Wu, C.T., & Woldorff, M.G. (2011). Sandwich masking eliminates both visual awareness of faces and face-specific brain activity through a feedforward mechanism. J. Vis. 11, 3.

27. Harris, J.A., McMahon, A.R., & Woldorff, M.G. (2013). Disruption of visual awareness during the attentional blink is reflected by selective disruption of late-stage neural processing. J. Cogn. Neurosci. 25, 1863–1874.

28. Hatamimajoumerd, E., Ratan Murty, N.A., Pitts, M.A., & Cohen, M.A. (2022). Decoding perceptual awareness across the brain with a no-report fMRI masking paradigm. Curr. Biol. 32, 4139–4149.

29. Kapoor, V., Dwarakanath, A., Safavi, S. Werner, J., Besserve, M., Panagiotaropoulos, T.I., & Logothetis, N.K. (2022). Decoding internally generated transitions of conscious contents in the prefrontal cortex without subjective reports. Nat. Commun. 13, 1535.

30. King, J.R., Faugeras, F., Gramfort, A., Schurger, A., El Karoui, I., Sitt, J.D., Rohaut, B., Wacongne, C., Labyt, E., Bekinschtein, T., Cohen, L., Naccache, L., & Dehaene, S. (2013). Single-trial decoding of auditory novelty responses facilitates the detection of residual consciousness. Neuroimage. 83, 726–738.

31. King, J.R., & Dehaene, S. (2014). Characterizing the dynamics of mental representations: the temporal generalization method. Trends Cogn. Sci. 18, 203–210.

32. Kleiner, M., Brainard, D., & Pelli, D. (2007). What’s new in Psychtoolbox-3. Perception. 36, 1–16.

33. Koch, C., Massimini, M., Boly, M., & Tononi, G. (2016). Neural correlates of consciousness: progress and problems. Nat. Rev. Neurosci. 17, 307–322.

34. Koivisto, M., & Revonsuo, A. (2007). Electrophysiological correlates of visual consciousness and selective attention. Neuroreport. 18, 753–756.

35. Kouider, S., Stahlhut, C., Gelskov, S.V., Barbosa, L.S., Dutat, M., de Gardelle, V., Cristophe, A., Dehaene, S., & Dehaene-Lambertz, G. (2013). A neural marker of perceptual consciousness in infants. Science. 340, 376–380.

36. Kroenmer, S.I., Aksen, M., Ding, J.Z., Ryu, J.H., Xin, Q., Ding, Z., Prince, J.S., Kwon, H., Khalaf, A., Forman, S., Jin, D.S., Wang, K., Chen, K., Hu, C., Agarwal, A., Wafa, S.M.A., Morgan, O.P., Wu, J., Christison-Lagay, K.L., Hasulak, N., Morrell, M., Urban, A., Constable, R.T., Pitts, M., Richardson, R.M., Crowley, M.J., & Blumenfeld, H. Human visual consciousness involves large scale cortical and subcortical networks independent of task report and eye movement activity. Nat. Comm., 13, 7342.

37. Lamme, V.A.F. (2018). Challenges for theories of consciousness: seeing or knowing, the missing ingredient and how to deal with panpsychism. Philos. Trans. R. Soc. B. 373, 1755.

38. Marti, S., & Dehaene, S. (2017). Discrete and continuous mechanisms of temporal selection in rapid visual streams. Nat. Commun. 8, 1955.

39. Mashour, G.A., Roelfsma, P., Changeux, J.P., & Dehaene, S. (2020). Conscious processing and the global neuronal workspace hypothesis. Neuron. 105, 776–798.

40. Naccache, L. (2018). Why and how access consciousness can account for phenomenal consciousness. Philos. Trans. R. Soc. B. 373, 1755

41. Nasman, V.T. & Rosenfeld, J.P. (1992) Taking attention to task: P300, Task response probability, and within-category deviation detection. Psychophysiol. 29, 657–663.

42. Navajas, J., Ahmadi, M., & Quiroga, R.Q. (2013). Uncovering the mechanisms of conscious face perception: a single-trial study of the N170 responses. J. Neurosci. 33, 1337–1343.

43. Odegaard, B., Knight, R.T., & Lau, H. (2017). Should a few null findings falsify prefrontal theories of conscious perception? J. Neurosci. 37, 9593–9602.

44. Paßler, M. (2023) The exclusionary approach to consciousness. Neurosci. Conscious. 10.1093/nc/niad022.

45. Pitts, M.A., Martinez, A., & Hillyard, S. (2012). Visual processing of contour patterns under conditions of inattentional blindness. J. Cogn. Neurosci. 24, 287–303.

46. Pitts, M.A., Padwal, J., Fennelly, D., Martínez, A., & Hillyard, S.A. (2014). Gamma-band activity and the P3 reflect post-perceptual processes, not visual awareness. Neuroimage. 101, 337–350.

47. Polich, J. (2020) 50+ years of P300: Where are we now? Psychophysiol. E13616.

48. Reiss, J.E. & Hoffman, J.E. (2007). Disruption of early face recognition processes by object substitution masking. Vis. Cogn. 15, 789–798.

49. Rodríguez, V., Thompson, R., Stokes, M., Brett, M., Alvarez, I., Valdes-Sosa, M., & Duncan, J. (2012). Absence of face specific cortical activity in the complete absence of awareness: converging evidence from functional magnetic resonance imaging and event-related potentials. J. Cogn. Neurosci. 24, 396–415.

50. Romei, V., Brodbeck, V., Michel, C., Amedi, A., Pascual-Leone, A., & Thut, G. (2008). Spontaneous fluctuations in posterior alpha-band EEG activity reflect variability in excitability of human visual areas. Cereb Cortex. 18, 2010–2018.

51. Salti, M., Monto, S., Charles, L., King, J.R., Parkkonen, L., & Dehaene, S. (2015). Distinct cortical codes and temporal dynamics for conscious and unconscious percepts. eLife. 4, e05652.

52. Sandberg, K., Bahrami, B., Kanai, R., Barnes, G.R., Overgaard, M., & Rees, G. (2013). Early visual responses predict conscious face perception within and between subjects during binocular rivalry. J. Cogn. Neurosci. 25, 969–985.

53. Schurger, A., Sarigiannidis, I., Naccache, L., Sitt, J.D., & Dehaene, S. (2015). Cortical activity is more stable when sensory stimuli are consciously perceived. Proc. Natl. Acad. Sci. USA. 112, E2083–E2092.

54. Sergent, C., Baillet, S., & Dehaene, S. (2005). Timing of the brain events underlying access to consciousness during the attentional blink. Nat. Neurosci. 8, 1391–1400.

55. Sergent, C., Corazzol, M., Labouret, G., Stockard, F., Wexler, M., King, J.R., Meyniel, F., & Fressnitzer, D. (2021). Bifurcation in brain dynamics reveals a signature of conscious processing independent of report. Nat. Commun. 12, 1149.

56. Shafto, J., & Pitts, M.A. (2015). Neural signatures of conscious face perception in an inattentional blindness paradigm. J. Neurosci. 35, 10940–10948.

57. Squires, K.C., Squires, N.K., & Hillyard, S.A. (1975) Decision-related cortical potentials during an auditory signal detection task with cued observation intervals. J. Exp. Psychol.: Hum. Percept. Perform. 1, 268–279.

58. Suzuki, M. & Noguchi, Y. (2013). Reversal of the face-inversion effect in N170 under unconscious visual processing. Neuropsychologia. 51, 400–409.

59. Tagliabue, C.F., Veniero, D., Benwell, C.S.Y., Cecere, R., Savazzi, S., & Thut, G. (2019) The EEG signature of sensory evidence accumulation during decision formation closely tracks subjective perceptual experience. Sci. Rep. 9(1):4949.

60. Tsuchiya, N., Wilke, M., Frässle, S., & Lamme, V.A.F. (2015). No-report paradigms: extracting the true neural correlates of consciousness. Trends Cogn. Sci. 19, 757–770.

61. Verleger, R. (2020). Effects of relevance and response frequency on P3b amplitudes: Review of findings and comparison of hypotheses about the process reflected by P3b. Psychophysiol. E13542.

62. Watson, A.B., & Pelli, D.G. (1983). Quest: a Bayesian adaptive psychometric method. Percept. Psychophys. 33, 113–120.

63. Womelsdorf, T., & Fries, P. (2007). The role of neuronal synchronization in selective attention. Curr. Opin. Neurobiol. 17, 154–160.

64. Woolgar, A., Duncan, J., Manes, F., & Fedorenko, E. (2018). Fluid intelligence is supported by the multiple-demand system not the language system. Nat. Hum. Behav. 2, 200–204.

65. Ye, M., Lyu, Y., Sclodnick, B., & Sun, H-J. (2019). The P3 reflects awareness and can be modulated by confidence. Front. Neurosci. 13, 519.

